# Single-cell and spatial multiomic inference of gene regulatory networks using SCRIPro

**DOI:** 10.1101/2023.12.21.572934

**Authors:** Zhanhe Chang, Yunfan Xu, Xin Dong, Yawei Gao, Chenfei Wang

## Abstract

The accurate reconstruction of gene regulation networks (GRNs) from sparse and noisy single-cell or spatial multi-omics data remains a challenge. Here, we present SCRIPro, a comprehensive computational framework that robustly infers GRNs for both single-cell and spatial multi-omics data. SCRIPro first addresses sample sparseness by a density clustering approach. SCRIPro assesses transcriptional regulator (TR) importance through chromatin reconstruction and *in silico* deletion, referencing 1,292 human and 994 mouse TRs. It combines TR-target importance scores with expression levels for precise GRN reconstruction. Finally, we benchmarked SCRIPro on diverse datasets, it outperforms existing motif-based methods and accurately reconstructs cell type-specific, stage-specific, and region-specific GRNs.

## Background

Transcription regulators (TR), including transcription factors (TF) and chromatin regulators (CR), play a crucial role in gene regulation by influencing transcription rates through mechanisms like recruiting transcriptional initiating complexes and modulating chromatin accessibility[1]. TRs form complex gene regulation networks (GRNs) with their target genes, also known as regulons, which are highly dynamic in different cellular contexts and serve as the foundation for various biological processes. Traditional methods for inferring GRNs, such as GENIE3[2], LISA[3], GRNBoost2[4], TIGRESS[5], ppcor[6], and NIMEFI[7], were primarily designed for bulk samples from mixed cell type tissues. However, these approaches are limited in their ability to accurately capture the regulatory programs operating in different cell types and states. With the advent of single-cell technologies, there has been a surge in the development of methods for inferring GRNs from single-cell transcriptome or epigenome data, including SCENIC[8], PIDC[9], SCODE[10], SINCERITIES[11], chromVAR[12], and SCRIP[13].

However, methods that solely focus on single-cell transcriptome for predictions not only neglect the genuine chromatin accessibility state but also fail to achieve single-cell resolution in GRN predictions, only attaining a cluster level. Additionally, many of these methods heavily rely on motif references to identify potential targets, which results in the loss of the cell type specificity and cannot robustly predict TR activity without motifs, particularly for chromatin regulators. To address these limitations, we previously developed SCRIP[13], a method that reconstructs single-cell TR activity and GRNs from scATAC-seq data by integrating extensive collections of TR ChIP-seq and motif references. Nonetheless, SCRIP’s effectiveness is influenced by the universality and the quality of scATAC-seq data. Recent advancements in single-cell multi-omics data have led to the development of new tools for predicting TR activity. For example, tools such as FigR[14], GRaNIE[15], DIRECT-NET[16] and GLUE[17] utilize paired or integrated multiome data as inputs and employ linear/non-linear regression methods to construct gene regulatory networks. Other tools, like CellOracle[18], offer pre-built GRNs or the ability to create custom-defined GRNs using scATAC-seq data. Meanwhile, tools like MICA[19] use bulk ATAC-seq to identify potential TR binding sites, applying this landscape to refine single-cell transcriptomic data [20]. However, it is worth noting that the aforementioned methods still heavily depend on the motif information, and further exploration is required to enhance their accuracy and coverage.

The flourishing development of spatial omics enables precise analysis of cellular structures within complex tissues and the spatial interactions between cells. Techniques such as Stereo-seq[21] and STARmap plus[22], have achieved single-cell-level resolution. These methods have significantly enhanced our understanding of gene regulation in specific microenvironments, providing valuable insights into crosstalk between microenvironment interactions and intracellular GRNs. However, most existing tools for predicting TR activity do not consider cellular spatial location. They overlook the impact in expression similarity of different cells or spots within the same microenvironment, leading to inaccurate TR predictions based on spatial transcriptomics data. Furthermore, the emergence of spatial multi-omics technologies, such as spatial ATAC-RNA-seq[23], provides paired chromatin accessibility states with gene expression. This presents an opportunity for accurate TR activity and GRN prediction using spatial multiomic information.

In this study, we have developed SCRIPro, a computational framework designed to predict TR activity and reconstruct TR-centered GRNs for both single-cell and spatial multiomic data. SCRIPro addresses the challenge of sparse single-cell or spatial multiomic signals by employing a density clustering approach that considers either expression or spatial similarities. Additionally, SCRIPro leverages a comprehensive TR reference compiled from TR ChIP-seq peaks obtained from Cistrome DB[24], along with motifs for 1,252 human TRs and 994 mouse TRs. Finally, SCRIPro combines TR-target importance from epigenomic data with TR-target expression from transcriptomic data to construct the GRNs. We demonstrate the robustness and versatility of SCRIPro by applying it to various datasets, including human B-cell lymphoma, mouse hair follicle development single-cell multi-ome data, mouse embryo Stereo-seq datasets at consecutive developmental time points, and P22 mouse brain Spatial-ATAC-RNA data. The results showcase the superior performance and utility of SCRIPro in diverse biological contexts.

## Results

### SCRIPro combines comprehensive chromatin and TR binding references to predict GRNs for both single-cell and spatial multi-omics data

SCRIPro is composed of two modules that can be applied to transcriptome and paired multiomic data. The expression module takes single-cell or spatial transcriptome as input. To address the limitations of gene coverage and minimize drop-out effects, SCRIPro employs a density clustering approach based on K-nearest neighbor (KNN) and a graph attention auto-encoder to generate SuperCells[25] with similar expression patterns or spatial coordinates[26] (Fig. 1). The expression profiles of SuperCells are then used to evaluate TR expression levels and co-expression patterns between TR and targets. The chromatin module of SCRIPro utilizes single-cell or spatial ATAC-seq as input. For transcriptome-only data, SCRIPro curates a comprehensive collection of public chromatin references encompassing 1,471 DNase-seq data and 2,575 H3K27ac data for both human and mouse samples[24] (Fig S1A-F). Subsequently, SCRIPro employs a logistic regression-based approach to scan the chromatin reference and reconstruct *in silico* chromatin landscapes that best match marker genes identified in SuperCells (Fig S1A, see Methods).

**Fig. 1.**
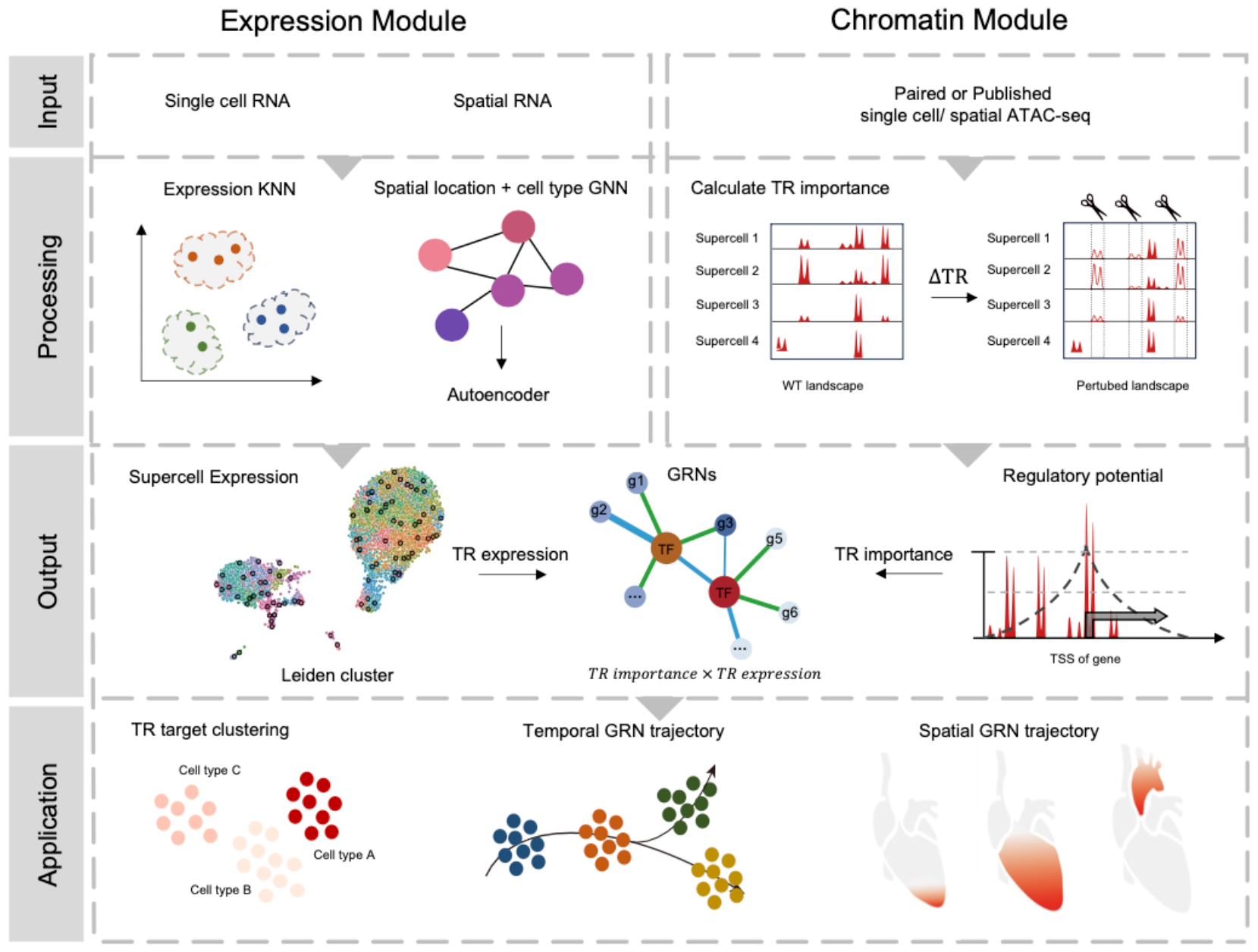
Overview of SCRIPro. SCRIPro takes single cell RNA-seq or spatial RNA-seq as input. SCRIPro first employs density clustering using a high coverage SuperCell strategy. While for spatial data, SCRIPro combines gene expression and cell spatial similarity information to a latent low-dimension embeddings via a graph attention auto-encoder. Then SCRIPro conducts *in silico* deletion analyses, utilizing matched scATAC-seq or reconstructed chromatin landscapes from public chromatin accessibility data, to assess the regulatory significance of TRs by RP model in each SuperCell. At last, SCRIPro combines TR expression and TR to generate TR-centered GRNs at the SuperCell resolution. The output of SCRIPro can be applied for TR target clustering, temporal GRN trajectory and spatial GRN trajectory.

SCRIPro leverages the paired or *in silico* reconstructed chromatin landscapes to assess the importance of TRs. Initially, SCRIPro complies a comprehensive TR reference dataset comprising 2,314 human TR ChIP-seq data and 1,920 mouse TR ChIP-seq data (Fig. S2A-B, Table S1). Additionally, this reference dataset is augmented with high-quality motifs obtained from cis-BP and HOMER, resulting in an extensive coverage of 1,252 human TRs and 994 mouse TRs (Fig S2C-F). SCRIPro then conducts *in silico* deletion analyses of TR binding sites to evaluate the impact of TRs on the expression of marker genes within SuperCells (Fig. 1 and Fig. S1A, see Methods). The potential targets of a TR are assigned based on the best-matched TR ChIP-seq data for a given SuperCell using a regulatory potential (RP) model (Fig. S1A). Finally, SCRIPro integrates the TR-target expression from transcriptome data and TR-target importance from epigenome data to generate the final TR activity within a SuperCell, identify the regulated genes, and construct the TR-centered GRNs (Fig. 1, Fig. S2G-J). The performance of SCRIPro is further assessed through downstream benchmarking, including accuracy evaluation of TR activity, clustering analysis, identification of cell type or stage-specific GRNs, and region-specific GRNs across diverse biological systems.

### SCRIPro identifies tumor-specific GRNs in the human B-cell lymphoma 10X multi-ome dataset

We first benchmarked the performance of SCRIPro on a human B-cell lymphoma (small lymphocytic lymphoma, SLL) dataset using 10X single-cell multi-ome containing 14,566 cells. We annotated 9 major cell types based on the expressed marker genes of each lineage, and tumor B-cells form a slightly different cluster compared to normal B-cells (Fig. 2A). SCRIPro accurately predicts well-known master regulators including SPI1, IRF1, FLI1, and STAT2 for mono/macrophages, GATA3, RUNX3, SMARCA4, JUND, and MYB for T-cells, PAX5 and IRF4 for B and tumor B-cells (Fig. 2B). The identified TR ChIP-seq reference corresponds well to the chromatin landscape from scATAC-seq at the SuperCell level, suggesting that the ChIP-seq reference is informative in predicting TR binding events (Fig. 2C). We compared the performance of SCRIPro with SCENIC+, a widely used algorithm that infers GRNs based on a combination of motif enrichment and GRNBoost2[4]. SCRIPro successfully identifies IRF8 in plasmacytoid dendritic cells (pDCs), which has been reported to be essential for the development of pDC and type 1 conventional dendritic cells[27]. However, SCENIC+ fails to assign IRF8 scores, possibly due to the limited number of pDC cells. SCRIPro specifically enriches SPIB, a driver regulator that mediates apoptosis through the PI3K-AKT pathway in diffuse B-cell lymphoma[28], in tumor B-cells but not normal cells, while SCENIC+ fails to predict the SPIB activity in all the B-cells. Finally, we assume the inferred TR activity should be highly correlated with their gene expression, with a positive correlation for activators and a negative correlation for repressors. SCRIPro shows a significantly high concordance between TR activity and TR expression compared to SCENIC+, with over 80% (81/105) of the factors showing better consistency (Fig. 2E, Fig. S3A-C Importantly, SCRIPro predicts activity scores for nearly 800 TRs, while SCENIC+ is only able to evaluate over 100 TR activities (Fig. S3B). In summary, these analyses suggest that SCRIPro can accurately infer GRNs globally and outperforms existing methods in terms of consistency between TR activity and TR expression.

**Fig. 2.**
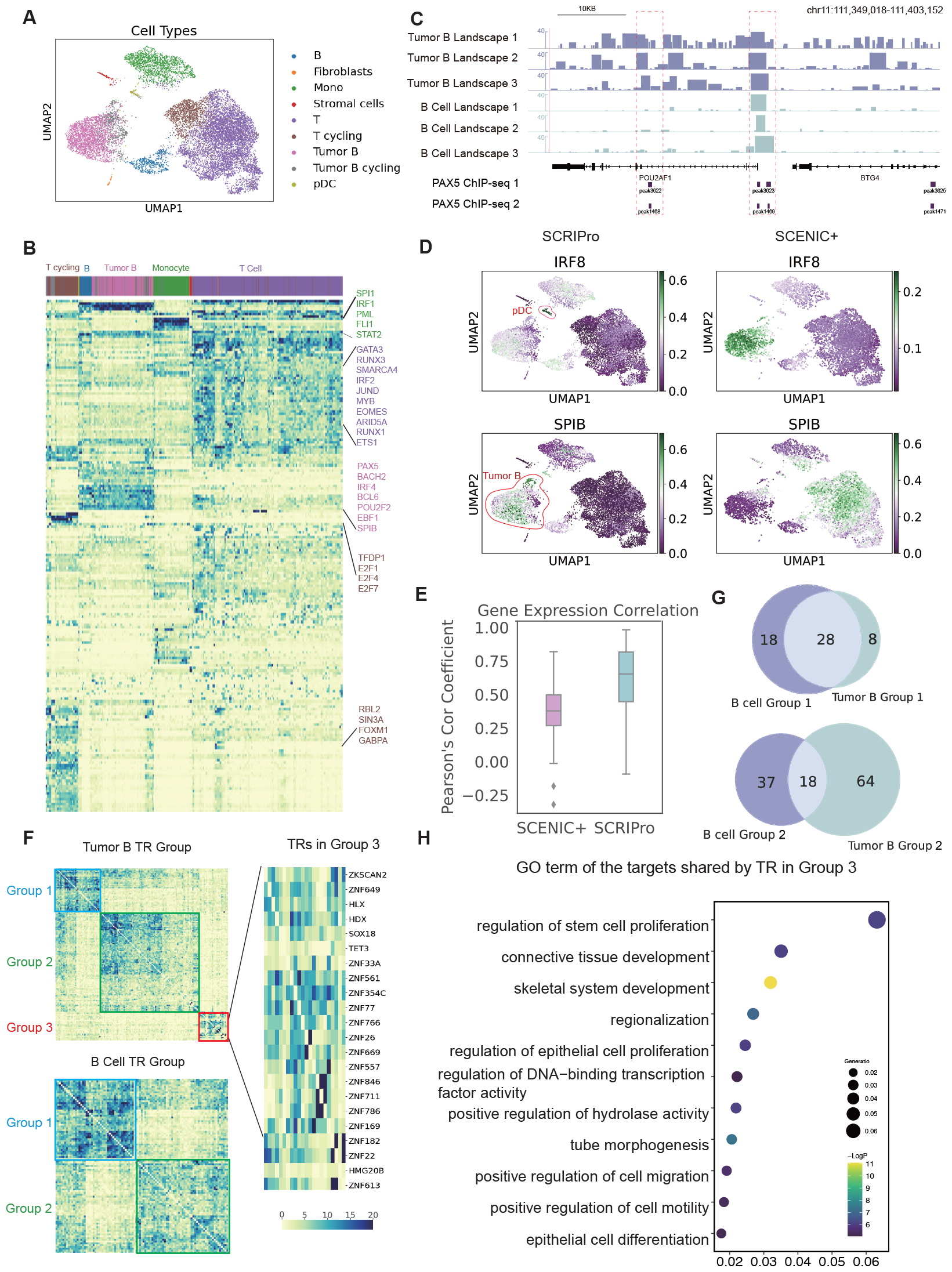
SCRIPro identified tumor-specific GRNs in the human B-cell lymphoma 10X multi-ome dataset. A. UMAP of 9 cell types identified in human B-cell lymphoma dataset. Mono: Monocytes. pDC: Plasmacytoid dendritic cells. B. Heatmap showing the clustering of TRs by cell type. Top: Cell types annotated in Figure A. Right: Highlighted TRs. C. The PAX5 ChIP-seq signal landscape identified by SCRIPro, which is aggregated based on supercells, show across tumor B and B cell types for POU5F1 and BTG4 genes on chr11:111349018-111403152. D. UMAP showing the predicted distributions of IRF8 and SPIB by SCRIPro and SCENIC+. IRF8 is highlighted in the pDC cell type in SCRIPro, while SPIB is prominent in Tumor B cells. E. Box plot depicting the Pearson correlation between TRs and gene expression by SCRIPro and SCENIC+. F. Heatmap clustering of TRs in tumor B and B cell types on the SCRIPro TR activity scale. Top: TR heatmap in the tumor B cell type, showing 3 clusters. Bottom: TR heatmap in the B cell type, showing 2 clusters. Right: An enlarged view of the group 3 (outlined in red) of the heatmap for tumor B, with all TRs labeled on the right. G. Top: Venn diagram showing the overlap of TRs between B cell group 1 and tumor B cell group 1. Bottom: Venn diagram showing the overlap of TRs between B cell group 2 and tumor B cell group 2. H. GO terms of the targets shared by TRs in Group 3.

B-cell lymphoma develops as a result of abnormal interactions between B cells and the microenvironment during development[29]. Given the accurate identification of TR activities specific to tumor B-cells, including SPIB, using SCRIPro, we conducted a systematic analysis to identify tumor-specific GRNs that potentially drive malignancy. Unsupervised clustering of TR activity identifies three independent clusters in tumor B-cells, compared to two clusters in normal B-cells (Fig. 2F). Both Group 1 and 2 were shared between tumor and normal B-cells, with Group 1 enriched in B-cell activation and differentiation, and Group 2 enriched in cytoplasmic translation that is important for B-cell development (Fig. 2G, Fig. S4A-D). Notably, the majority of TRs in Group 3 of tumor B-cells belonged to the ZNF family, which has been reported to silence retrotransposons and regulate epithelial proliferation[30, 31] (Fig. 2F-H). These analyses suggest that tumor B-cells may employ alternative proliferation strategies, such as activating epithelial proliferating genes. Collectively, our analyses demonstrate the superior accuracy and sensitivity of SCRIPro in identifying cell-type-specific TRs, even enabling the discrimination between similar cell types.

### SCRIPro reveals epigenetic priming effects of mouse hair follicle differentiation

TRs play a crucial role in driving cell type differentiation, compared to static datasets that only contain differentiated cells, reconstructing GRNs from developmental datasets presents challenges due to subtle differences along the development trajectories. We next benchmarked the performance of SCRIPro on a hair follicle differentiation dataset generated using the SHARE-seq protocol[32]. The cells were annotated into 7 major cell types including outer root sheath (ORS), transit-amplifying cells-1 (TAC-1), and TAC-2, medulla, hair shaft-cuticle cortex (cortex), inner root sheath (IRS), and mix cells (Fig. 3A). Starting from ORS cells, pseudo time analyses suggest three distinct differentiation paths that led to the formation of medulla, cortex and IRS cells (Fig. 3B). SCRIPro robustly identifies TRs enriched in the initial ORS cell type and the three different trajectories (Fig. 3C, Fig. S5 and Fig. S6A). For instance, medulla cells exhibited unique TR activity including Prdm1, Pbx3, and Rnf2, known for their significant regulatory roles in medullary-related functions[33] [34]. Cortex cells were characterized by specific activities of Lef1[35] and Rora[36]. Furthermore, IRS cells exhibited high activity of Gata3, an important TR in the skin stem cell lineage during the initiation of epidermal stratification and hair follicle IRS patterning[37] (Fig. 3D). These analyses demonstrate the capability of SCRIPro to identify lineage-specific GRNs along the developmental trajectory even with subtle differences.

**Fig. 3.**
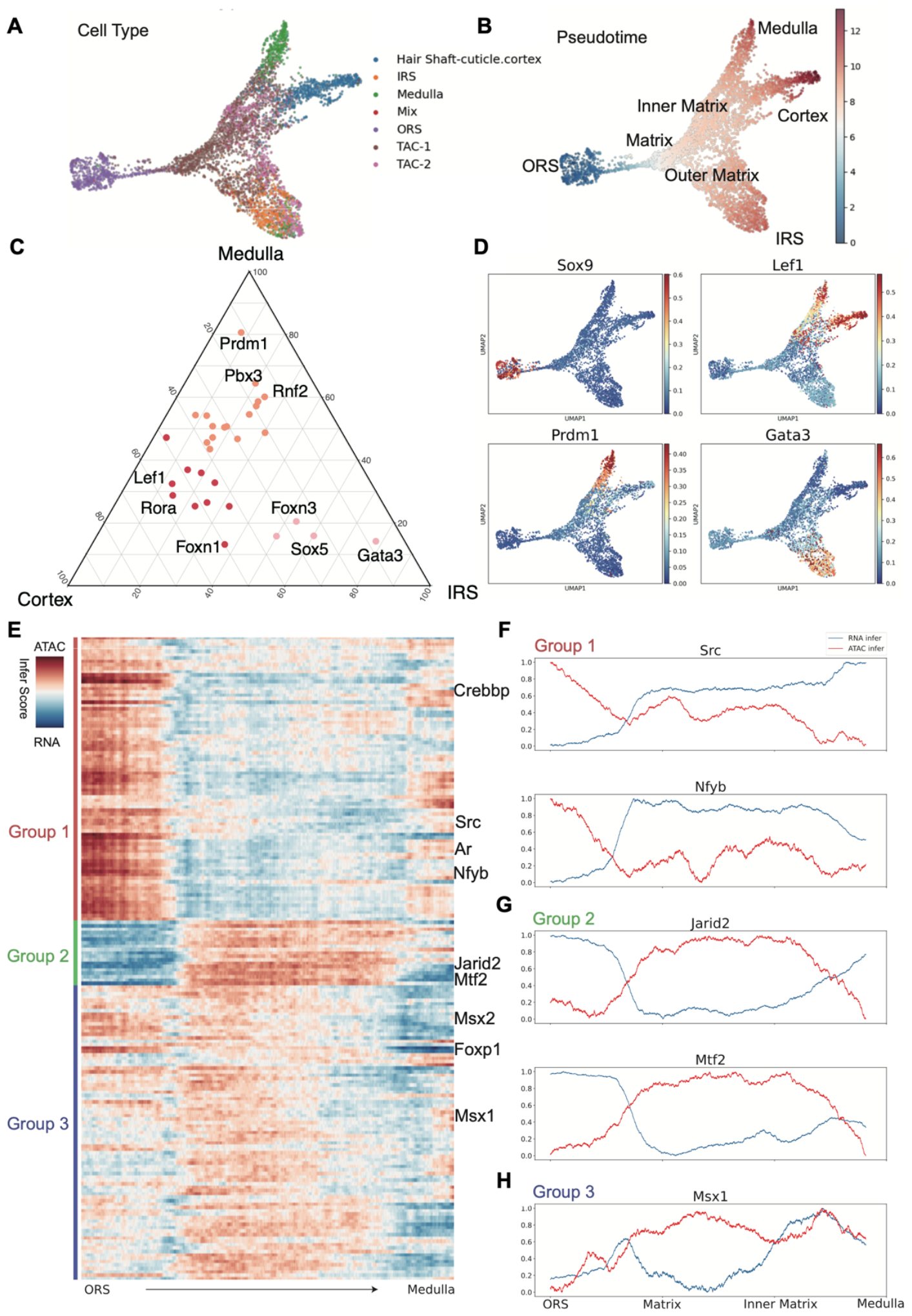
SCRIPro reveals epigenetic priming effect in the mouse hair follicle development SHARE-seq data. A. UMAP showing 7 cell types during hair follicle development. IRS: Inner Root Sheath. ORS: Outer Root Sheath. B. Pseudotime UMAP plot illustrating the developmental trajectory of hair follicle structure, with different nodes representing various tissue types. C. Ternary plot showing TRs enriched in Medulla, Cortex, and IRS tissue types, respectively. Colors represent different tissue types, and numbers indicate the relative enrichment level. D. UMAP showing the distribution of TR activity of Sox9, Lef1, Prdm1, and Gata3. E. Heatmap illustrating the clustering of all TRs along the developmental trajectory from ORS to medulla, based on the difference in ATAC and RNA values. Left: 3 groups identified by the heatmap clustering. Infer score: The difference in rank values of all TRs between SCRIP and SCRIPro. Red indicates SCRIP - SCRIPro > 0, while blue indicates SCRIP – SCRIPro < 0. F-H. RNA infer score and ATAC infer score of group 1-3 (Src, Nfyb, Jarid2, Mtf2 and Msx1) from ORS to medulla.

Traditional GRN prediction methods designed for scRNA-seq datasets, such as SCENIC, PIDC, and SCODE, primarily rely on the co-expression information to infer TR regulation. However, the prediction of TR activity based on gene expression and chromatin accessibility can be decoupled due to the influence of epigenetic priming. Since SCRIPro could both reconstruct the TR activity from expression (transcriptome-only module) and chromatin accessibility (epigenome-only or paired module), we compared the difference between RNA versus ATAC inferred TR activity to systematically evaluate the potential priming effect. We focused on specific developmental lineage, for example, the ORS to medulla path. Encouragingly, our analysis revealed a TR group in which chromatin-inferred activity preceded expression-inferred activity (Fig. 3E-F, Group 1), indicating a strong epigenetic priming effect that leads to a delay in target gene expression. Conversely, Group 2 exhibited an opposite trend, with RNA-inferred activity preceding ATAC-inferred activity (Fig. 3G). Notably, many factors in this group possess repressive functions including Jarid2 and Mtf2, subunits of the PRC2 complex reported in mouse embryonic stem cells[38], thus validating the priming effect of repressive H3K27me3 modifications. Group 3, comprising the majority of TRs, displayed largely synchronous patterns regardless of whether they were predicted using RNA or ATAC data (Fig. 3E, H). Similar patterns were observed for the ORS to cortex and ORS to IRS directions (Fig. S6B). In summary, these analyses confirm the epigenetic priming effect using the hair follicle development SHARE-seq data and also underscore the importance of chromatin landscapes in accurately predicting the TRs required for the differentiation of future trajectories.

### Integration of spatial information enhances TR activity and GRN prediction on E16.5 mouse embryo Stereo-seq data

To showcase the superior performance of SCRIPro in reconstructing spatial GRNs by integrating spatial locations and neighborhood information, we applied it to a Stereo-seq dataset derived from E16.5 mouse embryos[21] (Fig. 4A). Our spatial clustering strategy successfully identified 28 cell types with unique spatial locations, and exhibiting denser cell positioning and higher within-cell type homogeneity compared to the traditional KNN-based method using Scanpy (Fig. 4A, Fig. S7A). We next quantified the TR activity in different cell types and re-clustered the cells by TR activity. SCRIPro demonstrated a significantly higher consistency with the cell types generated by spatial clustering compared to SCENIC (Fig. S7B-C). For instance, we examined the TRs Foxo1 and Foxo3, known to be enriched in the brain and facial regions including the striatum, anterior thalamic nucleus, olfaction, and dental epithelium[39]. SCRIPro robustly predicted the TR activity in these regions, with Foxo3 exhibiting a dispersed distribution compared to Foxo1[39]. In contrast, SCENIC failed to enrich either factor in the brain and facial regions (Fig. S7D). Gata6 plays a critical role in cardiac function with pronounced enrichment in the outflow tract and aortic arch[40], SCRIPro distinctly identified the Gata6 activity across different spatial subsets of the heart (Fig. S7D). Additionally, SCRIPro exhibited a broader TR coverage and successfully predicted the spatial TR activity for Neurod2 and Otx2 in distinct brain regions, which were not covered by SCENIC (Fig. S7E). Finally, we benchmarked the performance of SCRIPro on homologous factors with similar motifs. SCRIPro accurately localized Gata4 in the heart region and Gata1 in the developing liver region, despite the highly resemblant motifs of these two factors (Fig. 4B). Similarly, SCRIPro demonstrated the ability to distinguish between Tcf12 and Tcf7, demonstrating the ability in identifying cell type-specific binding patterns of homologous factors through incorporating ChIP-seq data (Fig. S7F). Collectively, these results suggest SCRIPro could accurately predict cell type and region-specific GRNs than existing tools on the spatial transcriptomic-only dataset, with the ability to distinguish TRs with similar motif patterns.

**Fig. 4.**
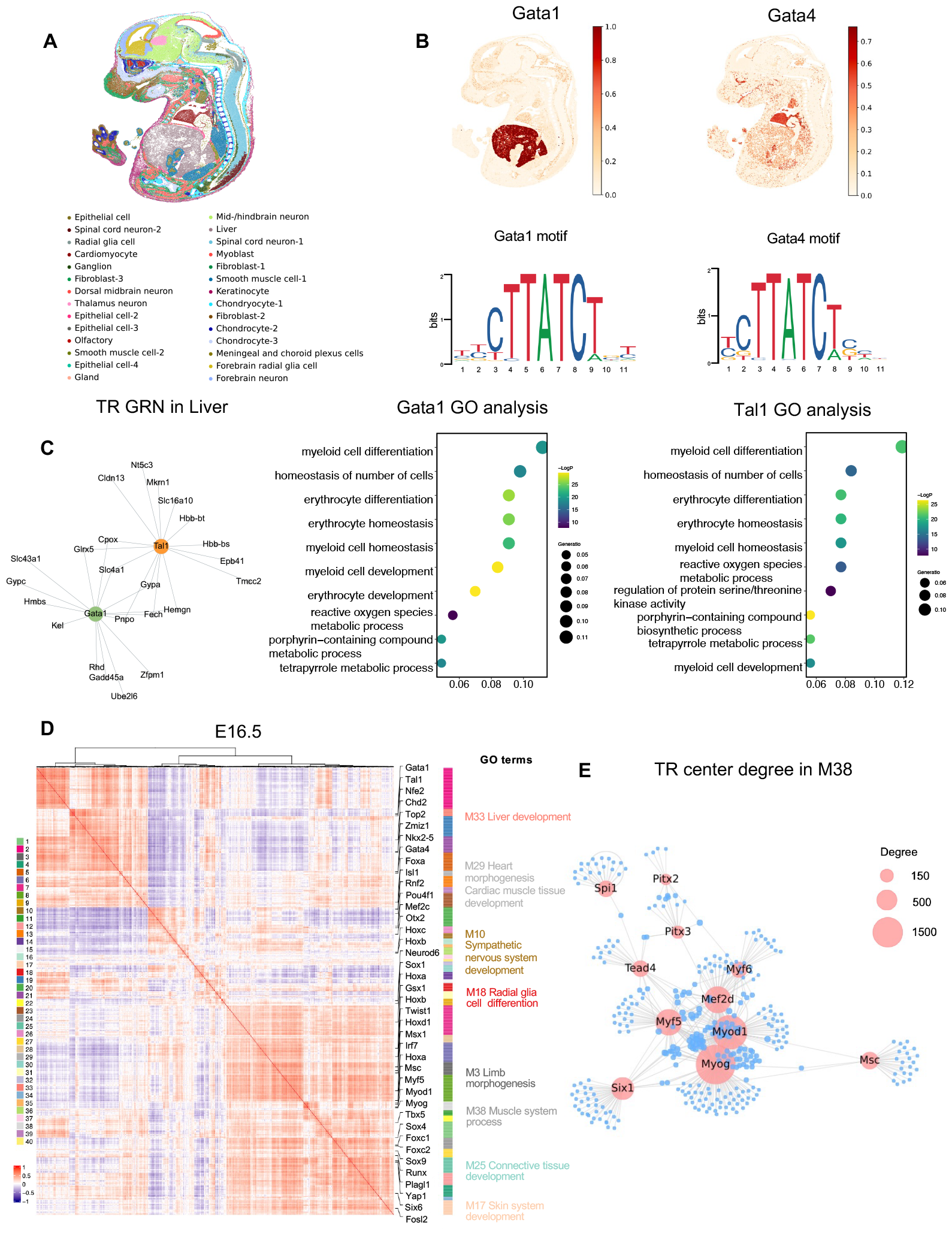
SCRIPro detected cell type specific GRNs in E16.5 mouse embryo Stereo-seq data. A. SCRIPro identified 28 cell types on E16.5 mouse embryo stereo-seq dataset. B. SCRIPro can distinguish different TRs spatial distribution with similar motif in same family. C. SCRIPro is capable of predicting Gata1 and Tal1 target genes and build GRNs, and utilizes these target genes for GO analysis. D. Heatmap showing the modules with significant spatial autocorrelation that are clustered into different modules based on spatial co-expression of E16.5 embryo. E. Calculate TR center degree in module 38 (muscle) and find out important TRs.

TRs and their associated cofactors collaborate to modulate downstream genes, thereby establishing GRNs instrumental in determining cell phenotypes. To evaluate the effectiveness of SCRIPro in identifying TR regulons, we focused on factors enriched in the embryonic liver, specifically Gata1[41] and Tal1[42]. We conducted a screening of their target genes and pruned the network based on TR-target co-expression, resulting in the generation of co-regulation regulons (Fig. 4C). Functional analyses revealed that both TRs are linked to myeloid cell and erythrocyte differentiation and homeostasis, indicating a potential co-binding of these two TRs in regulating hematopoietic function in the embryonic liver (Fig. 4C). Similarly, by iteratively clustering the spatially variable TRs and regulon co-expression patterns, we successfully identified 40 co-regulated gene modules in the mouse E16.5 embryo (Fig. 4D and Fig. S8A-B). These modules show highly specific spatial patterns, including Pax5 for forebrain (M1), Neurog2 for mid/hindbrain (M10), Myog for muscle cells (M38), Msx1 in maxillary and limb mesenchymal cells[43] (M3), and Hoxc8 in the mouse embryonic spine[44] (M36) (Fig. 4D and Fig. S8B). Within each module, SCRIPro also constructed TR-centered GRNs to identify crucial factors. For example, Myod1, Myog, Mef2d, and Myf5 were identified as key nodes in the M38 muscle module, aligning well with their known roles as muscle master regulators (Fig. 4E). In summary, these analyses demonstrate the ability of SCRIPro to accurately identify TR target genes and construct cell type-specific GRNs, thereby enabling the identification of potential novel master regulators for each lineage.

### SCRIPro detects stage-specific GRNs in cardiomyocytes across mouse embryonic heart sections

Our previous analyses demonstrate that SCRIPro could identify lineage- and stage-specific GRNs in the hair follicle development dataset, we next evaluated whether it is suitable for analyzing time-series spatial datasets. We applied SCRIPro on mouse embryo stereo-seq data[21] spanning from E11.5 to E15.5 across five continuous stages (Fig. 5), and aligned these spatial slides using SLAT[45] (Fig. 5A). Our focus was on heart development, given its early formation in mammalian embryos and the diverse changes observed in cardiomyocytes from E11.5 to E15.5 (Fig. 5B). Cells in the heart region were annotated into fibroblasts (FBs, marked by Postn), epicardium (marked by Ptn), and cardiomyocytes (CM, marked by Ttn) at E11.5, and CMs further differentiated into ventricle cardiomyocytes (vCMs, marked by Myh7) and atrial cardiomyocytes (aCMs, marked by Myl7) at E13.5 and E15.5 (Fig. S9A-B). Encouragingly, SCRIPro identifies highly specific TR activity among different cell types starting from E11.5, including Gata6, Jund, and Mef2c for cardiomyocytes, Tcf4 and Hoxb3 for epicardium, and Sox9 for fibroblasts (Fig. 5C). Interestingly, at E13.5, there was still a significant overlap in the TRs of aCMs and vCMs, with Tbx5 enriched in both lineages, Tbx20 and Tbx18 specifically enriched in aCMs, and Mef2d enriched in vCMs.

**Fig. 5.**
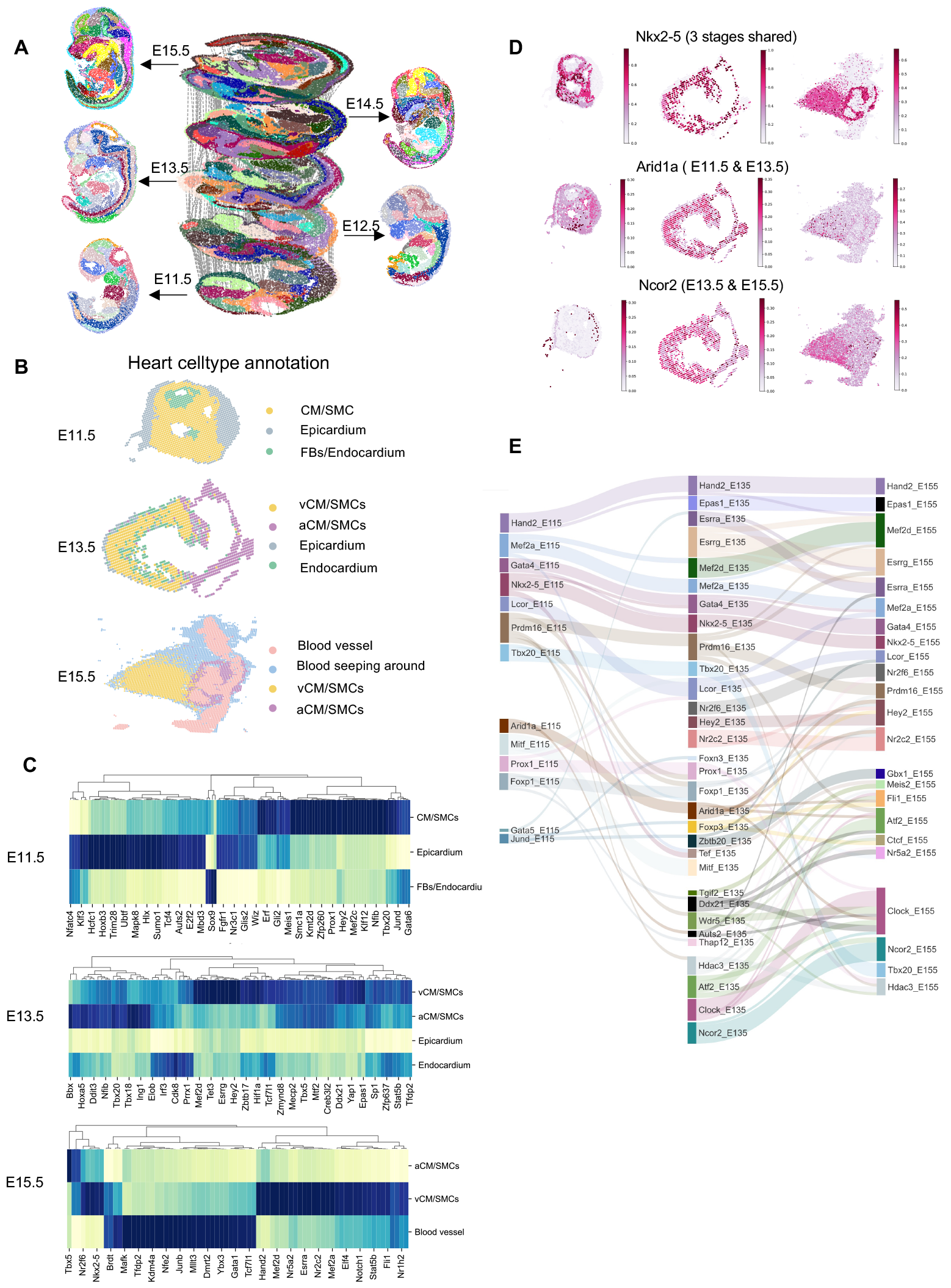
SCRIPro identified stage-specific GRNs in consecutive mouse embryonic heart sections. A. Alignment of embryonic sections from five developmental stages, E11.5, E12.5, E13.5, E14.5, and E15.5, based on clusters predicted by SCRIPro. B. Annotation of embryonic hearts at three developmental stages: E11.5, E13.5, and E15.5. C. Highly specific TR activity among different cell types starting from E11.5 to E15.5. D. TR spatial distribution in 3 stages. Nkx2-5: 3stages shared. Arid1a: E11.5 and E13.5 shared. Ncor2: E13.5 and E15.5 shared. E. The stage specific GRNs correlation across three developmental stages.

However, at E15.5, most of the TRs showed remarkable differences, with Tbx5 only enriched in aCMs and Mef2d still enriched in vCMs (Fig. 5C and Fig. S9C). In summary, our analyses revealed that Mef2d might play an important role in vCM differentiation at the E13.5 stage, while Tbx5 functions later to regulate aCM differentiation at E15.5.

To further elucidate the dynamics of TRs and their regulons during cardiomyocyte development, we constructed a cross-stage GRN based on enriched TRs in cardiomyocytes (Fig. S9D). Our analyses identified stage-shared regulators including Nkx2-5, Hand2, Gata4, and Mef2d, as well as early-stage regulators Arid1A and late-stage regulator Ncor2. Most of these factors are supported by existing literatures[46, 47] (Fig. 5D and Fig. S9D). In addition to TR activity, SCRIPro robustly identified the TR regulons, for which can be used to evaluate the stable or dynamic regulation of different TRs (Fig. 5E). Most TRs showed conserved regulation of its regulon among different stages. However, Prdm16, a regulator for brown adipocyte differentiation[48], shows remarkable dynamics in its regulon[49, 50] (Fig. 5E). We performed functional analyses of different subsets of the Prdm16 regulon and found that the constant Prdm16 regulon was enriched in fatty acid beta-oxidation, consistent with its well-known regulatory functions (Fig. S10). Interestingly, the genes lost at E11.5 were highly enriched in glycogen metabolism, while the unique genes at E13.5 were specifically enriched in NADP metabolism (Fig. S10). These analyses suggest that Prdm16 may play a critical role in the metabolic reprogramming from glycolysis to oxidative phosphorylation in cardiomyocytes, which has been reported to be connected with the proliferation ability of cardiomyocytes[51, 52]. In summary, SCRIPro effectively tracks and analyzes the dynamics of TRs as well as their regulons in mouse embryonic heart development, enabling future identification of novel lineage regulators from spatial transcriptomics-only datasets.

### Spatial multi-omic prediction of GRNs reveals crosstalk between intra-cellular gene regulation and extra-cellular interactions in the P22 mouse brain

Finally, we applied SCRIPro to a spatial ATAC-RNA-seq dataset from the mouse brain on postnatal day 22, which provided paired expression and chromatin accessibility data along with spatial location. SCRIPro successfully identified 10 distinct spatial domains that corresponded well with specific anatomical structures (Fig. 6A). Notably, domains 0, 1, and 3 represented Cortex (CT) regions associated with advanced neural and emotional processing functions[53]. Domain 6 corresponded to the Corpus Callosum (CC) region, crucial for interhemispheric communication, while domain 9 aligned with the Lateral Ventricle (LV) region, known for neural stem cell origins. Pseudo-time analysis revealed an inward-to-outward neural stem cell emergence pattern consistent with the differentiation trajectories of neuron cells (Fig. S11A). SCRIPro accurately predicted spatial variable TRs for each region. For instance, it identified Sox6 and Sox10 as significant TRs in the CC region, and Sox2, Sox4, and Sox11 in the LV regions, highlighting the pivotal role of the SOX family in neurogenesis[54] (Fig. 6B). Additionally, Bcl11b and Foxp1[55] exhibit strong TR activity within the striatum, where Bcl11b plays a critical role in the differentiation of medium spiny neurons important to motor control[56] (Fig. 6B). Moreover, Mef2c[57], Neurod2, and Neurod6[58, 59] have been previously demonstrated to regulate gene expression in the CT region (Fig. 6C and Fig. S11B). These results validate the accuracy of SCRIPro in predicting region-specific TRs using sparse spatial multiomic datasets.

**Fig. 6.**
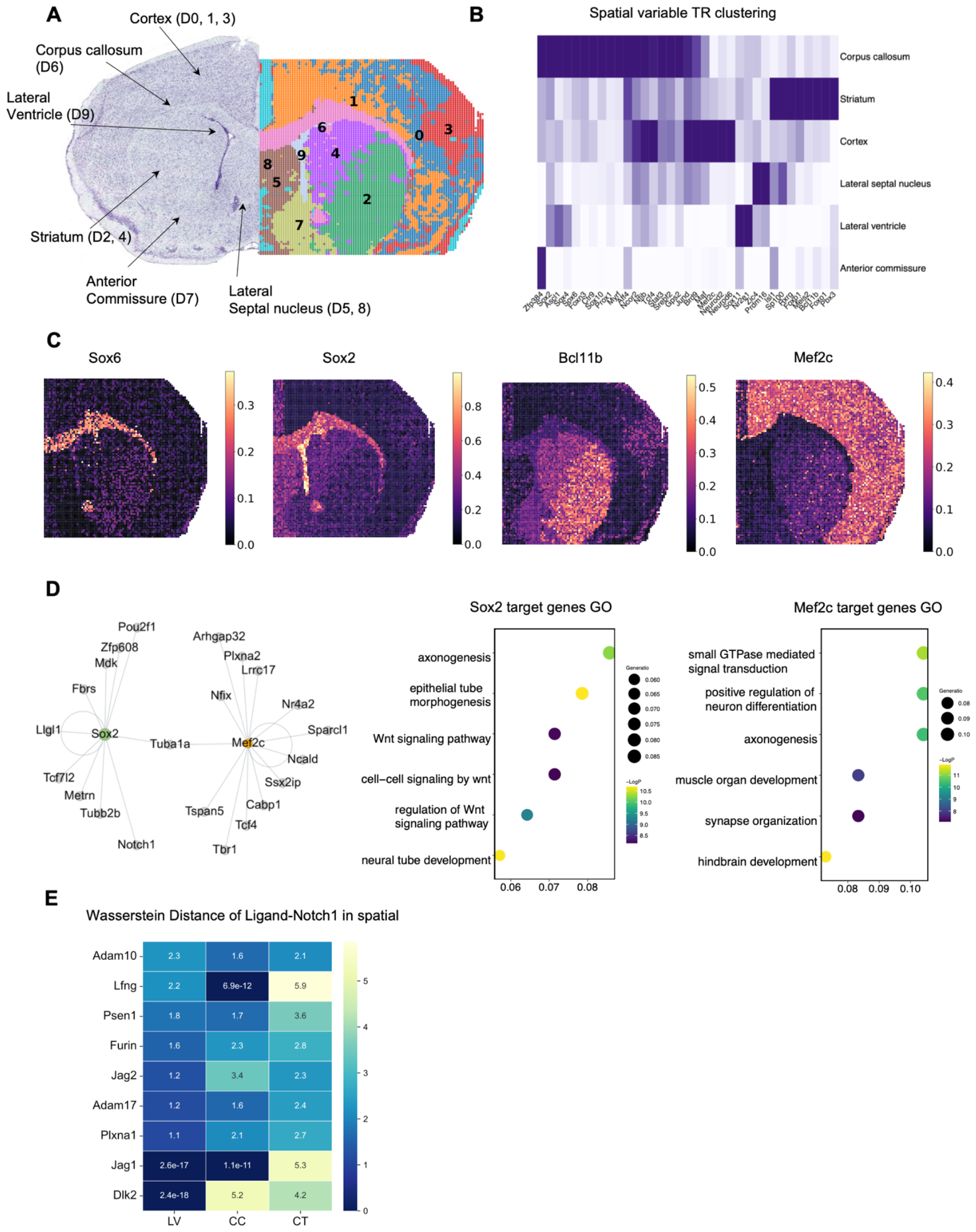
Spatial multi-omic prediction of GRNs reveals crosstalk between intra-cellular gene regulation and extra-cellular interactions in the P22 mouse brain. A. SCRIPro identified 10 clusters on mouse brain spatial ATAC-RNA-seq (RNA). B. Heatmap of spatial variable TR clustering by 6 cell type regions. C. SCRIPro activity score of selected marker TR in spatial. D. SCRIPro predicts Sox2 and Mef2c target genes then builds GRNs, and utilizes these target genes for GO analysis. E. Wasserstein distance of Ligand and Notch1 receptor in different regions.

To further assess SCRIPro’s accuracy in predicting TR regulons from spatial multiomic data, we constructed GRNs for Sox2 and Mef2c, which were enriched in LV and CT regions, respectively (Fig. 6D). The targets and functions of these two factors exhibited distinct characteristics. The Sox2 regulon was notably enriched in the WNT signaling pathway and neural tube/epithelial tube development[60], while the MEF2C regulon predominantly converged on signal transduction, synapse organization, and hindbrain development[61]. These analyses emphasize the specific functions of these TRs and demonstrate the accuracy of SCRIPro in identifying TR-specific regulons. In addition to intrinsic gene regulation, cellular crosstalk could also regulate cell type-specific TR expression. We then investigated whether spatial GRN analyses could be used to identify extracellular regulations on TRs such as cell-cell interactions. Specifically, we focused on the TRs with documented ligand-receptor pairs (see Methods). For example, we observed a high enrichment of NOTCH1 activity in the LV region (D9), which is known to harbor neural stem cells. Interestingly, its upstream regulator Dlk1/2[62] and Jagged1[63] were found to have closer associations with the NOTCH1 regulon in the LV region compared to the CC region (D6) and CT region (D0) (Fig. 6E). Similar analyses using FGFR1 expression did not yield significant differences (Supplementary Fig. S11C). In summary, SCRIPro effectively utilizes spatial multiomic data to construct detailed GRNs in the mouse brain, revealing distinct TR specificity across different spatial regions and facilitating the exploration of extracellular regulations influencing TR expression in different regions.

## Discussion

Constructing high-resolution GRNs from extensive single-cell and spatial transcriptomics data is crucial for understanding mechanisms of gene regulation in cell fate determination and disease development. In this study, we developed SCRIPro, a rapid and user-friendly method that accurately predicts GRNs. SCRIPro addresses the challenge of sparse signals at the SuperCell[25] resolution by taking account of expression and spatial similarity. Notably, SCRIPro includes a chromatin reconstruction step designed for scRNA-seq and spatial transcriptomics datasets without paired ATAC-seq data, significantly enhancing its usability. Additionally, through *in-silico* deletion TR analyses using a comprehensive TR ChIP-seq reference, SCRIPro outperforms existing motif-based methods in various tested systems, including human B-cell lymphoma, mouse hair follicle development, mouse developing embryos, and brain at P22. We anticipate that SCRIPro will be widely utilized by researchers to identify crucial TRs underlying novel differentiation and disease mechanisms.

Despite its current superiority over existing methods, SCRIPro still has some limitations. Firstly, its performance relies heavily on the quality of public ChIP-seq datasets, which can affect its robustness. To address this, SCRIPro integrates motif scanning results into the TR reference to mitigate the reduction in TR reference coverage after filtering out low-quality datasets. Furthermore, the rapid development of single-cell/spatial epigenomics technologies such as sciATAC-seq3[64], sciMAP-ATAC[65], and some multiome epigenomics technologies such as Paired-Tag[66], scCUT&Tag-pro[67] and DOGMA-seq[68], accelerates the accumulation of high-quality epigenome data in the field. SCRIPro aims to incorporate these data into the chromatin and TR reference, expanding coverage to more cell types and improving performance, particularly on rare cell types. Secondly, while SCRIPro adeptly predicts TRs at a SuperCell resolution, its precision at the single-cell level is still being established due to challenges related to background noise and data quality. To overcome these limitations, we are exploring the use of machine learning models, such as diffusion and autoencoders, to improve the quality of single-cell and spatial-omics data, thereby enhancing both the resolution and accuracy of TR prediction. Finally, the cellular spatial locations from spatial-omics data are essential for understanding *in situ* gene expression regulation, cell interactions, and signal transduction within spatial microenvironments. Currently, SCRIPro calculates upstream signaling pathways by constructing spatial GRNs and filtering ligand-receptor (L-R) pair expressions. However, this approach is limited by the coverage of spatial-omics data and the number of L-R pairs in the database. Future development integrating GRNs, protein-protein interactions (PPI), and L-R co-occurrence is expected to significantly increase the connections between intracellular GRNs and extracellular cell-cell interactions (CCIs).

In summary, SCRIPro is a promising tool that enables researchers to leverage extensive single-cell or spatial transcriptomics data to identify driver TR regulations, with or without paired epigenome information. It facilitates the interpretation of GRNs across diverse cell types, cellular trajectories, and spatial domains. These analyses will enhance our understanding of TR regulation in development systems, and the associations between TR dysfunction and disease occurrence.

## Methods

### Overview of the SCRIPro Method

To decrease computational requirements and to alleviate the impact of dropout events inherent in single-cell sequencing data, as well as to improve the stability of the inferred GRNs, we adopted a divide-and-conquer approach. SCRIPro begins by constructing Supercells[25], which are aggregates of gene expression profiles from clusters of individual cells exhibiting similar transcriptional activity. Following this initial step, SCRIPro identify a set of marker genes for each Supercell. These marker genes serve as representative features of their respective Supercells and are subsequently utilized as inputs for the LISA[3] framework. Within LISA, SCRIPro implement ISD analyses to assess the effects of transcriptional regulator (TR) perturbations on the expression of these marker genes. The outcome of this process is a GRN constructed at the Supercell level, which provides a higher-order representation of the regulatory landscape across the pooled cellular subpopulations.

### Reference Dataset

We employed the identical ChIP-seq datasets for SCRIP, which were sourced from the Cistrome Data Browser[24]. Following a meticulous reorganization and refinement of annotations pertaining to factors, cell types, and tissues, we applied stringent filtering criteria: a median quality score for raw sequences exceeding 25, a uniquely mapped reads proportion surpassing 50%, a PCR Bottleneck Coefficient (PBC) greater than 0.8, an enrichment of peaks by a minimum of 10-fold in quantities exceeding 100, a Fraction of Reads in Peaks (FRiP) above 0.01, and an overlap of the top 5000 peaks with joint DNase I hypersensitive sites (DHS) exceeding 70%. We preserved only those peaks demonstrating a minimum 5-fold enrichment within each dataset. Datasets comprising fewer than 1000 peaks were subsequently excluded. After this filtration process, we amassed a total of 2314 human and 1920 mouse TR ChIP-seq datasets, encompassing 671 and 440 TRs, respectively.

In an effort to enhance the representation of transcription factors, we also synthesized pseudo-peaks data by conducting motif scanning. We procured the transcription factor position weight matrix (PWM) motifs for both human and mouse, amalgamated and transformed the data formats, and subsequently utilized the HOMER[69] software to scan the genome for motif-associated genomic intervals. We juxtaposed the scanning outcomes with the Encyclopedia of DNA Elements (ENCODE)[70] candidate cis-regulatory elements (ccREs) and the Cistrome Union DHS compilation, excluded any intersections with known blacklisted regions, and augmented the length of each motif locus to 340 base pairs to facilitate comparison with the ChIP-seq datasets. By imposing a filter based on the P-value, we retained the top 25,000 binding sites, ultimately yielding 916 human and 816 mouse pseudo-peaks derived from motif scanning.

Subsequently, we integrated these meticulously curated SCRIPro datasets into the LISA framework, generating reference HDF5 (Hierarchical Data Format version 5) files for both human and mouse datasets, which comprised the count of ChIP-seq peaks and the metadata associated with the datasets. In summation, the constructed human TR index encompasses 1252 TRs, while the mouse TR index includes 997 TRs.

To assess the influence of transcriptional regulators on their target genes, we conducted a detailed analysis of ChIP-seq datasets contained in HDF5 files using the Regulatory Potential (RP) model[71]. The computational formula for RP, adhering to the SCRIP methodology, is as follows:

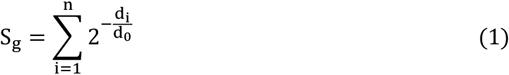

In this equation, n denotes the count of TR binding sites proximal to the TSS of gene *g*, while *d*_*i*_ signifies the distance from the i-th peak’s center to the TSS. For TRs exhibiting more than 20% of peaks in the promoter region, they are categorized as promoter-centric TRs, and the half-decay distance *d*_0_ is set at 1 kb. In contrast, TRs with fewer promoter-localized peaks are classified as enhancer-centric, with their half-decay distance established at 10 kb. To enhance computational efficiency, the analysis was restricted to genes located within a half-maximal regulatory range (half-decay distance) of 15 *d*_0_, as peak scores beyond this threshold diminish to less than 0.0005.

An enhanced version of the RP model was employed to explore the potential target genes regulated by transcription factors. This model extends beyond incorporating exon information by also accounting for the regulatory impact of adjacent genes. Specifically, if a peak is detected within the exon region of a gene, the corresponding score is assigned the value of 1, which is then normalized relative to the gene’s total exon length. Conversely, should a peak reside in the promoter or exon region of a neighboring gene, its score is designated as 0.

### Preprocessing

#### Single-cell RNA-seq data

We first preprocess counts matrix for each cell following Scanpy worflow[72]. Next, principal component analysis (PCA) is performed using *scanpy. tl. pca*() function to reduce data dimensionality. We compute neighboring cells for each cell using *scanpy. pp. neighbors*() function, setting the number of neighbors to N_neighbors (default 10) and the number of principal components to N_pcs (default 40). Finally, we apply *scanpy. tl. umap*() function for UMAP dimensionality reduction and perform Leiden clustering using *scanpy. tl. leiden*() function with a resolution parameter set to resolution (default 0.8) to stratify cellular populations.

#### Spatial transcriptomics RNA-seq data

After normalizing the count matrix, SCRIPro utilizes the approach from STAGATE[73] to build a spatial neighbor network (SNN), integrating the similarity between adjacent spots of a given location, and subsequently transforms this spatial data into an undirected neighbor network based on a predetermined radius *r*. Subsequently, utilizing a cell type-aware module, the SNN is pruned based on pre-clustered gene expression. After constructing SNN, SCRIPro employs a graph attention auto-encoder to integrate gene expression and spatial location. In cell type-aware module, SCRIPro employs a self-attention mechanism for both types of SNNs. The learned spatial similarities from the standard SNN and the cell type-aware SNN are denoted as **att**^*spatial*^ and **att**^*aware*^, respectively. The final spatial attribution used is a linear combination of these two (where α, the default hyperparameter set at 0.5, represents the weight of the cell type-aware SNN):

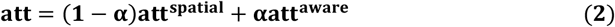

The output of the decoder is considered as the reconstructed normalized expressions. We then perform dimensionality reduction on the integrated data using *scanpy. pp. neighbors*() and *scanpy. tl. umap*(), following the same methodology as above.

#### Supercell Construction

After employing leiden clustering[74] to identify cell subsets at a resolution parameter of 0.8, each cluster is treated as an independent RNA-seq dataset. Using a binary search method, we iteratively increase the secondary resolution to obtain more refined leiden cluster classifications until the average number of cells in each small leiden cluster reaches a user-specified N. Supercells within each large Leiden cluster that contain fewer than 30 cells are merged with the nearest Supercell to ensure a minimum of 30 cells per Supercell.

Marker genes are then computed for each Supercell: For smaller scRNA-seq datasets, we identify the top 500 marker genes per Supercell using the *scanpy. get. rank*_*genes*_*groups*_*df*() method. For larger datasets (cell number > 150,000), we recommend you to use large-scale marker gene selection strategy. We extract top 1500 genes in each leiden cluster, and then identify genes expressed above the 60% percentile in each Supercell as marker genes for subsequent analyses. Supercells with fewer than 35 selected marker genes are excluded from further calculations.

#### GRN Inference

Upon identifying the marker genes for each Supercell, we input them into the LISA[3] framework. For scRNA-seq data, SCRIPro divides the input data into 8 chunks (by default) based on the number of parallel threads and performs LISA’s ISD calculations for each Supercell’s marker genes within the chunks, obtaining results for each Supercell. For multi-omics data, such as scMultiome-seq, which combines scRNA-seq and scATAC-seq data, we first align the datasets using GLUE[17] if the barcodes cannot be directly matched. If the barcodes are matchable, we aggregate the scATAC-seq data based on the Supercell division from the scRNA-seq data. To prepare the scATAC-seq data, we sort it using the bedtools[75] sort command and merge intervals within each sorted TSV file, ensuring that they do not exceed 1000 bases, using the bedtools merge command. Each merged TSV file is then converted into a bigwig format file using the *bedGraphToBigWig* command[76]. These bigwig files, corresponding to each Supercell, are used as the landscape during the ISD step.

LISA’s chromatin landscape model uses L1-regularized logistic regression to select an optimum sample set for H3K27ac ChIP-seq or DNase-seq samples. LISA first calculated the chrom-RP for each RefSeq gene. The chrom-RP for gene k in sample j is defined as

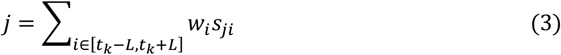

*L* is set to 100 kb, The weight *w*_*i*_ represents the regulatory impact of a locus at position *i* on the gene *k*’s transcription start site at genomic position *t*_*k*_. *s*_*ji*_ is the signal of chromatin profile j at position i.

LISA also calculates peak-RP for each set of ChIP-seq data. The definition of peak-RP is same to what has mentioned above. Then the ISD method recalculates the chrom-RP after erasing the signal in all 1-kb windows containing at least one peak from a putative regulatory cistrome, and then comparing the model RPs with and without deletion to produce a ΔRP value for each gene. The combined statistics method for TR ranking compares the peak-RPs or ΔRPs of the query gene set with that of the background gene set. It uses the one-sided Wilcoxon rank-sum test and combines peak-RP, DNase-seq, and H3K27ac chrom-RP for ChIP-seq-based methods. The Cauchy combination test is used to compute a summary p value for each TR. SCRIPro applies a negative logarithmic transformation to the summary p-values obtained from LISA’s ISD results, serving as the score for individual TRs within each Supercell.

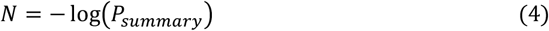

Additionally, for each TR and its corresponding targets within a Supercell, we calculate a z-score relative to the mean:

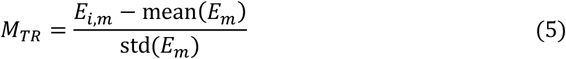

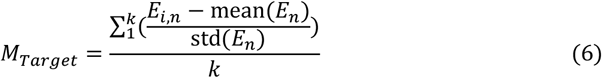

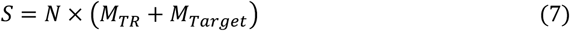

where E represents the expression value of TRs (represented by m) or target genes (represented by n), while i denotes each Supercell. More specifically, k represents the target genes regulated by TR m. For each set of ChIP-seq data, we select genes with an RP score > 5 as the target genes for this TF. If the number of genes with an RP score > 5 is less than 300, then we choose the top 300 genes in RP score ranking as the target genes.The individual TR’s score obtained in the previous step is multiplied by this value to derive the final TR activity score within that Supercell. For each TR and its corresponding target within a Supercell, we multiply their RP-score by the Pearson correlation coefficient between the TR and the target, which serves as the regulatory score for this TR-target pair within the Supercell.

### B-Cell Lymphoma Dataset

#### Preprocessing

In the scRNA-seq dataset, cells with fewer than 200 genes and genes present in fewer than 3 cells were removed. We retained only cells that had both RNA-seq and ATAC-seq counts. We applied SCRIPro with default parameters to the filtered scMultiome-seq peak count matrix to evaluate the activity of transcriptional regulators. The activity scores of different TRs were visualized as heatmaps with seaborn’s clustermap and projected onto UMAP plots with Scanpy. After integrating the original scATAC-seq data with SCRIPro, the resulting landscape was displayed using the IGV genome browser[77].

#### Clustering Performance Comparison

For the assessment of SCRIPro, we chose to compare it with SCENIC+[78]. The SCENIC algorithm[8], which is the precursor to SCENIC+, is a widely recognized method for GRN inference. SCENIC+ builds upon SCENIC by providing single-cell resolution transcription factor activity scores, allowing for a comprehensive performance comparison with SCRIPro.

Since SCRIPro relies on ChIP-seq datasets, it is not suitable for direct comparison with traditional methods that are based on synthetic GRN datasets. Instead, we compared SCRIPro with SCENIC+ in terms of gene expression correlation. Specifically, SCRIPro selects the TR activity score in Supercells and calculates the correlation with gene expression within those Supercells. On the other hand, SCENIC+’s results are correlated with the original cell expression. Both methods were used with their default parameters.

To visualize and cluster the number of target genes of TFs in Tumor B Cells and B Cells, we utilized the *sns. clustermap*() function in the Seaborn library, using the default parameters. Additionally, we employed Metascape[79] to calculate the GO terms of the overlapping genes identified by the two methods. Finally, we visualized the GO terms using an R script.

### Hair Follicle Development Dataset

#### Preprocessing

We annotated SHARE-seq data cell types using labels from the original study. SCRIPro was applied with default parameters to the SHARE-seq data to assess the activity of TRs in each cell.

#### Pseudotime Analysis

We utilized MIRA[80] for trajectory analysis of the SHARE-seq data. We designated ORS as the starting cell type and IRS, Cortex, and Medulla as the terminal cell types to study the differentiation trajectory in hair follicle development. Cells were ordered by pseudotime using the ‘mira_pseudotime’ column from MIRA results. The activity score of each TR in each terminal cell type (Medulla_prob/IRS_prob/Cortex_prob > 0.8) was visualized in ternary plots.

### ATAC-RNA Analysis

SCRIP[13] was used to perform TR activity analysis on the scATAC-seq data from the SHARE-seq data, applying the SCRIP enrichment function with default parameters to the peak count matrix. Subsequently, we transformed the numerical scores from two distinct matrices—single-cell RNA sequencing data (RNA-infer) and single-cell ATAC sequencing data (ATAC-infer)—into their corresponding ranks. We then proceeded to compute the discrepancies between the two matrices by subtracting the RNA-infer ranks from the ATAC-infer ranks, thereby deriving a set of differential rankings that culminate in final score. These scores were clustered and compared according to the pseudotime obtained from MIRA[80].

### Mouse Embryonic Development Dataset

#### Preprocessing

The Stereo-seq dataset detailing mouse embryonic development was procured from the specified database (https://db.cngb.org/search/project/CNP0001543/). SCRIPro build SNN and perform graph attention auto-encoder using STAGATE strategy. We utilized SCRIPro to process embryonic data at E16.5 to identify 28 spatial clusters employing a large dataset gene selection strategy. Standard preprocessing steps, including quality control, normalization, and data filtering, were executed prior to downstream analyses. Motif pattern was downloaded from JASPAR(https://jaspar.elixir.no/)[81].

#### Identification of TR Activity Co-expression Modules

The R package Giotto *binSpect*() function was employed to discern TRs exhibiting spatial coherence in their activity scores as determined by SCRIPro. Giotto[82] identified 40 distinct spatial TR modules in E16.5 embryo datatset, which were subsequently analyzed using heatmap clustering. Centrality metrics were calculated using networkx Python package *degree*_*centrality*() function.

#### TR Activity Analysis and Comparison

To assess the cluster purity of the identified spatial domains, we adopted ROGUE score (https://github.com/PaulingLiu/ROGUE). Transcription factor activity scores were computed using both SCENIC and SCRIPro, adhering to their default parameters. The re-clustering of these scores, integrating spatial context, facilitated the computation of the ROGUE score via the R package ROGUE. Normalized Mutual Information (NMI) scores were derived by comparing the clustering outcomes of TR activity scores obtained from both methods against those from STAGATE clustering.

### P22 Mouse Brain Spatial Multi-omics Dataset Analysis

#### Preprocessing

The spatial ATAC-RNA-seq multi-omics dataset of the P22 mouse brain was retrieved from the cited source (GSE205055). The *scripro. cal*_*ISD*_*parallel*() function from SCRIPro was then employed to analyze the multi-omics data, resulting in the determination of TR activity scores for each Supercell. We employed the *binSpect*() function in Giotto to identify spatially variable TRs and subsequently conducted heatmap clustering to discern cell type-specific TRs. The pseudotime spatial trajectory analysis of neural stem cell emergence was explored by spaceFlow[83].

#### Wasserstein distance calculation

To investigate the impact of signaling pathways on TR expression in spatial contexts, we used the Wasserstein distance[84] to measure differences in TR and gene expression across brain regions, utilizing spot positions as coordinates and expression levels as values. For a ligand expressed in *m* spots and a receptor expressed in *n* spots, we formed a matrix *D* ∈ *R*^*m*×*n*^ to record the Euclidean distances between spots, based on spatial coordinates. By identifying an optimal transport *γ*′ ∈ *R*^*m*×*n*^ that minimizes the total transport cost of ligand and receptor expression distributions: *L* ∈ *R*^*m*×1^ and *R* ∈ *R*^*n*×1^. This total transport cost is determined by summation of the products of the transport value and the Euclidean distance between each spot. Based on the optimal transport plan, the the Wasserstein distance can be computed as follows:

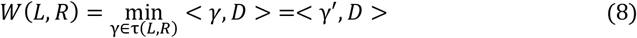

## Availability of data and materials

SCRIPro is now available at https://github.com/wanglabtongji/SCRIPro. All data used in this study have been previously reported and are publicly available (Supplementary Table S1). Single cell multiome ATAC-RNA-seq data is available from 10X genomics (https://www.10xgenomics.com/resources/datasets/fresh-frozen-lymph-node-with-b-cell-lymphoma-14-k-sorted-nuclei-1-standard-2-0-0) Hair follicle SHARE-seq data was downloaded from the GEO database (https://www.ncbi.nlm.nih.gov/geo/query/acc.cgi?acc=GSE140203). The mouse embryo is available from STOmicsDB MOSTA (https://db.cngb.org/stomics/mosta/download/). The mouse brain Spatial-RNA-ATAC-seq data was obtained from the GEO database (GSM6753043 spatial RNA-seq, GSM6758285 spatial ATAC-seq) by Fan lab.

## Acknowledgements

The authors thank the Bioinformatics Supercomputer Center of Tongji University for offering computing resource.

## Author information

Zhanhe Chang, Yunfan Xu and Xin Dong contributed equally to this work.

### Contributions

C.W. conceived and supervised the whole project. Y.X. and Z.C. designed and implemented the SCRIPro algorithm. X.D. collected and preprocessed the ChIP-seq and motif datasets. Z.C. evaluated the performance on mouse embryo stereo-seq datasets and mouse brain spatial ATAC-RNA-seq datasets. Y.X. performed the analysis of B-cell lymphoma 10X Genomics multiomic datasets and the hair follicle single-cell multiomic datasets. Z.C., Y.X., X.D., Y.G. and C.W. wrote the manuscript with the help of other authors. All authors read and approved the final manuscript.

### Corresponding authors

Correspondence to <u>Chenfei Wang</u>.

## Ethics declarations

### Ethics approval and consent to participate

Not applicable.

### Consent for publication

Not applicable.

## Supplementary Tables

**Supplementary Table 1** Datasets information used in this study.

**Supplementary Table 2** Summary of human TR reference datasets used in this study.

**Supplementary Table 3** Summary of mouse TR reference datasets used in this study.

