## Supplementary figures and images for "Single-cell and spatial multiomic inference of gene regulatory networks using SCRIPro"

### Supplementary figures

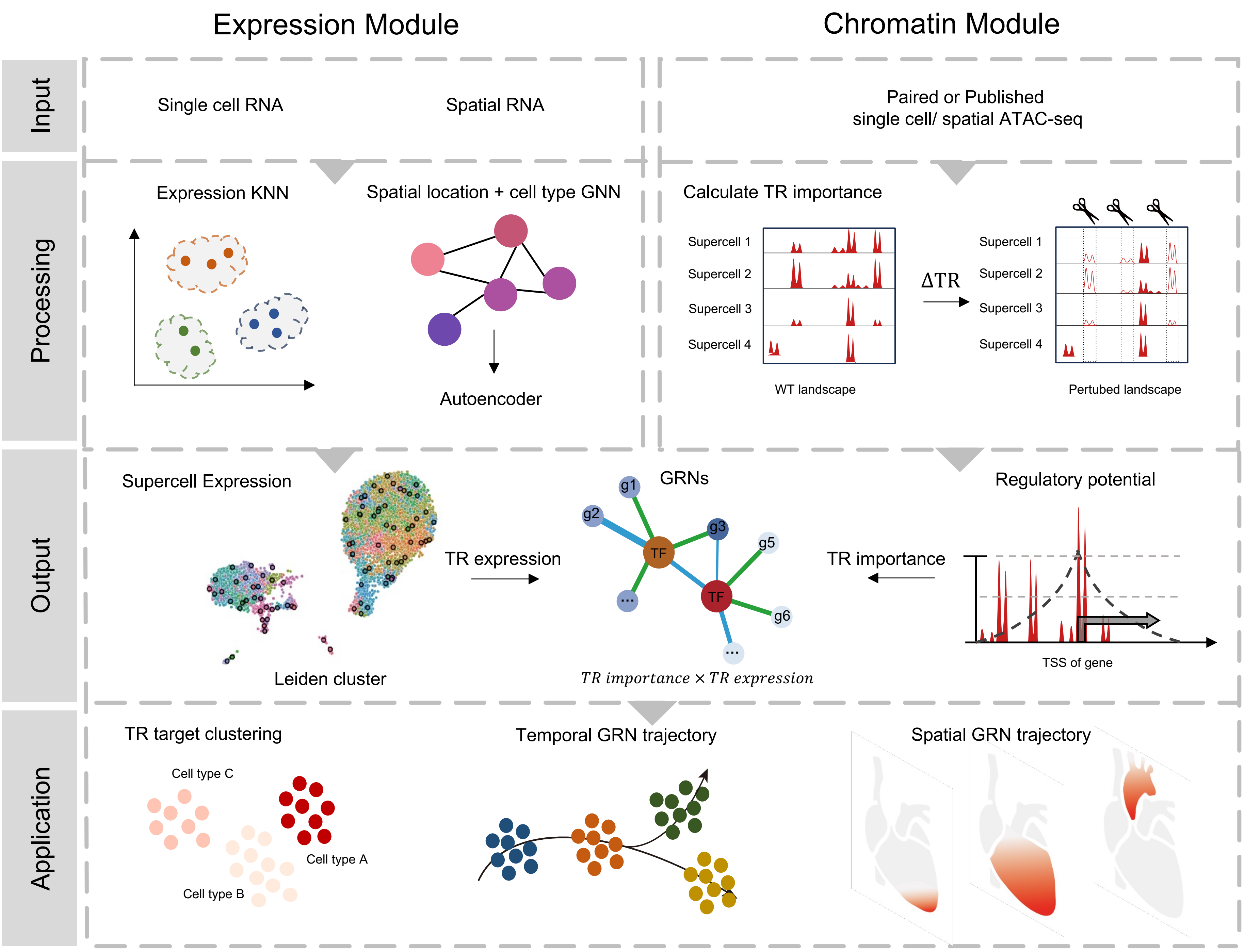

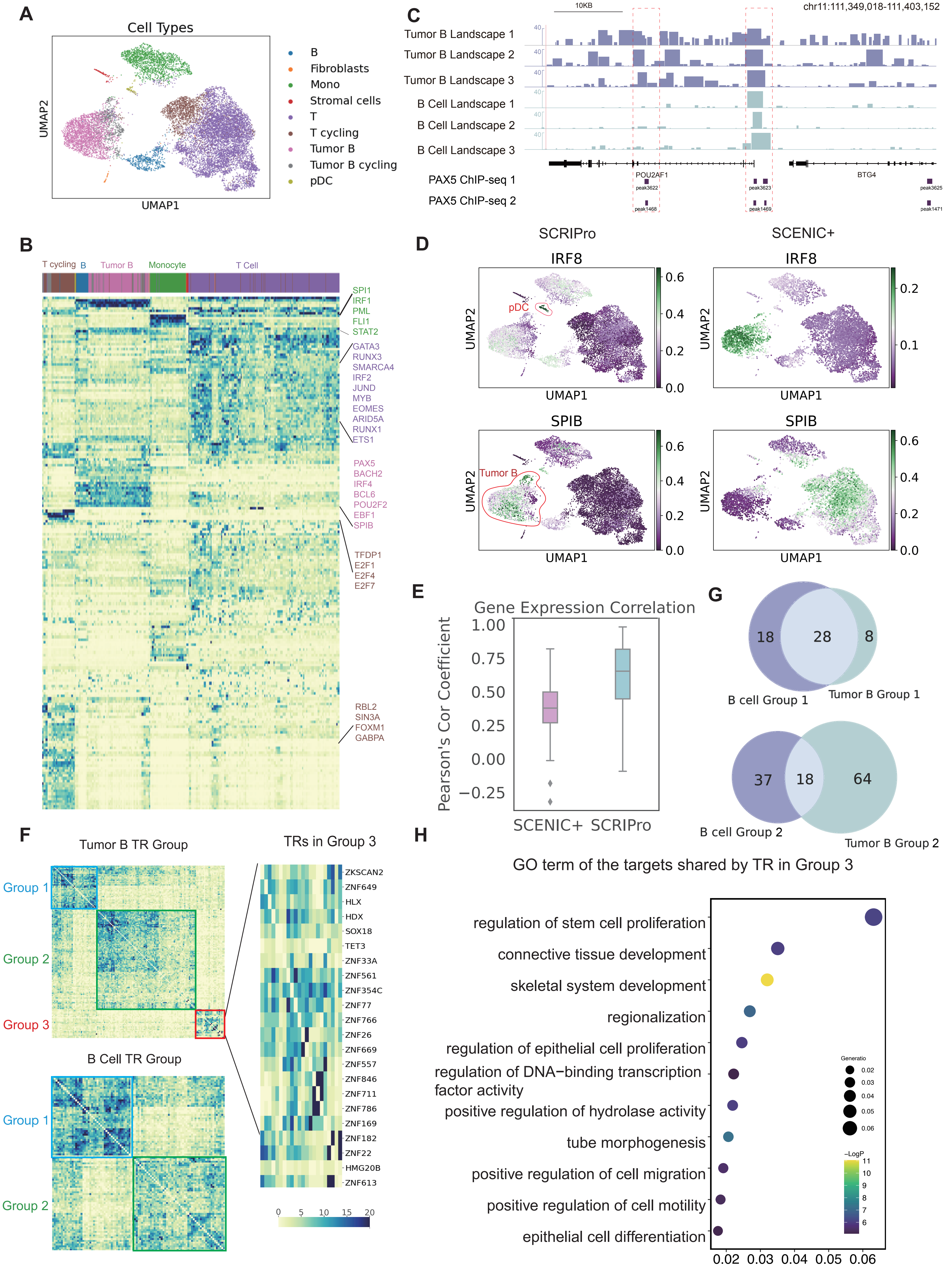

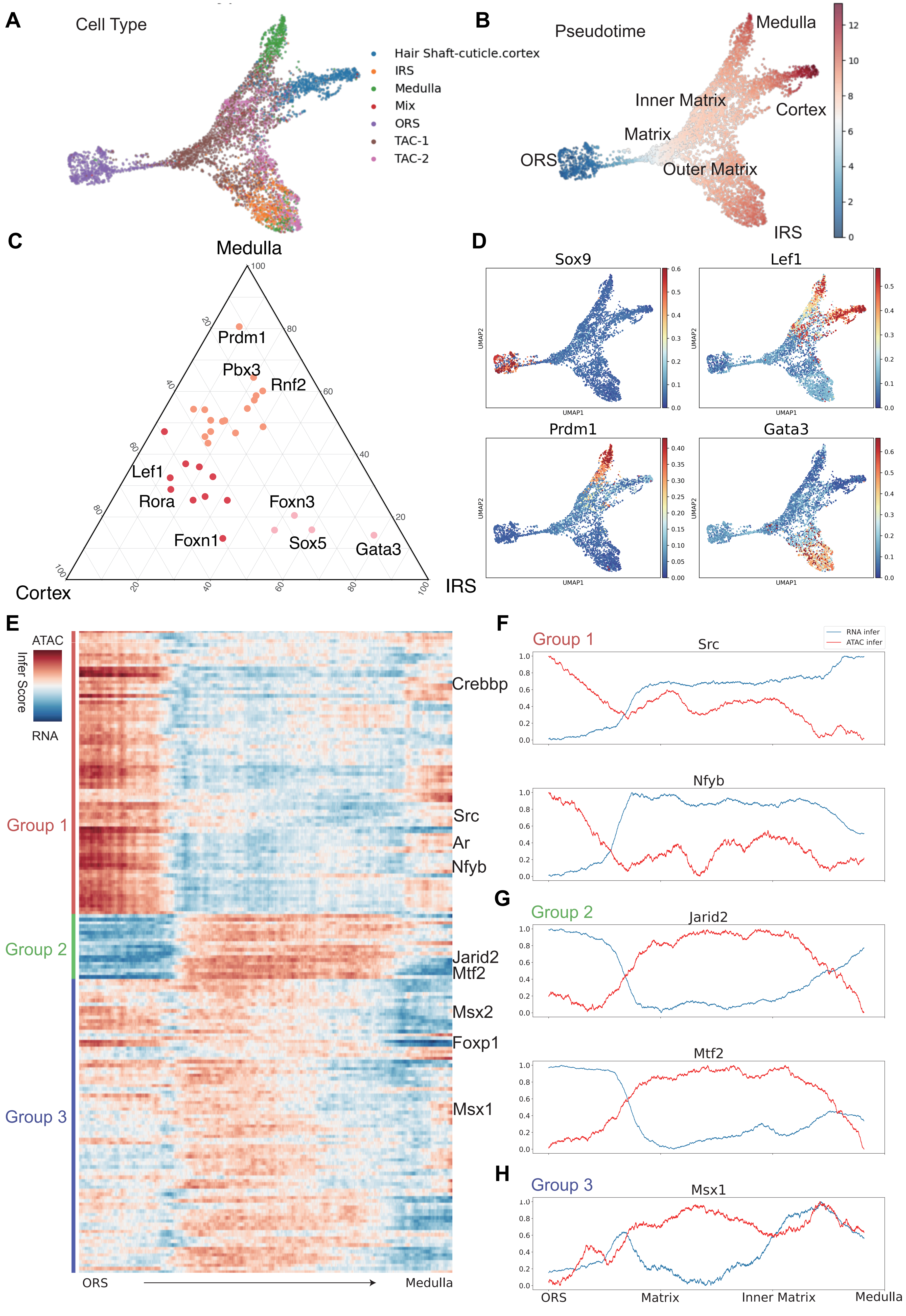

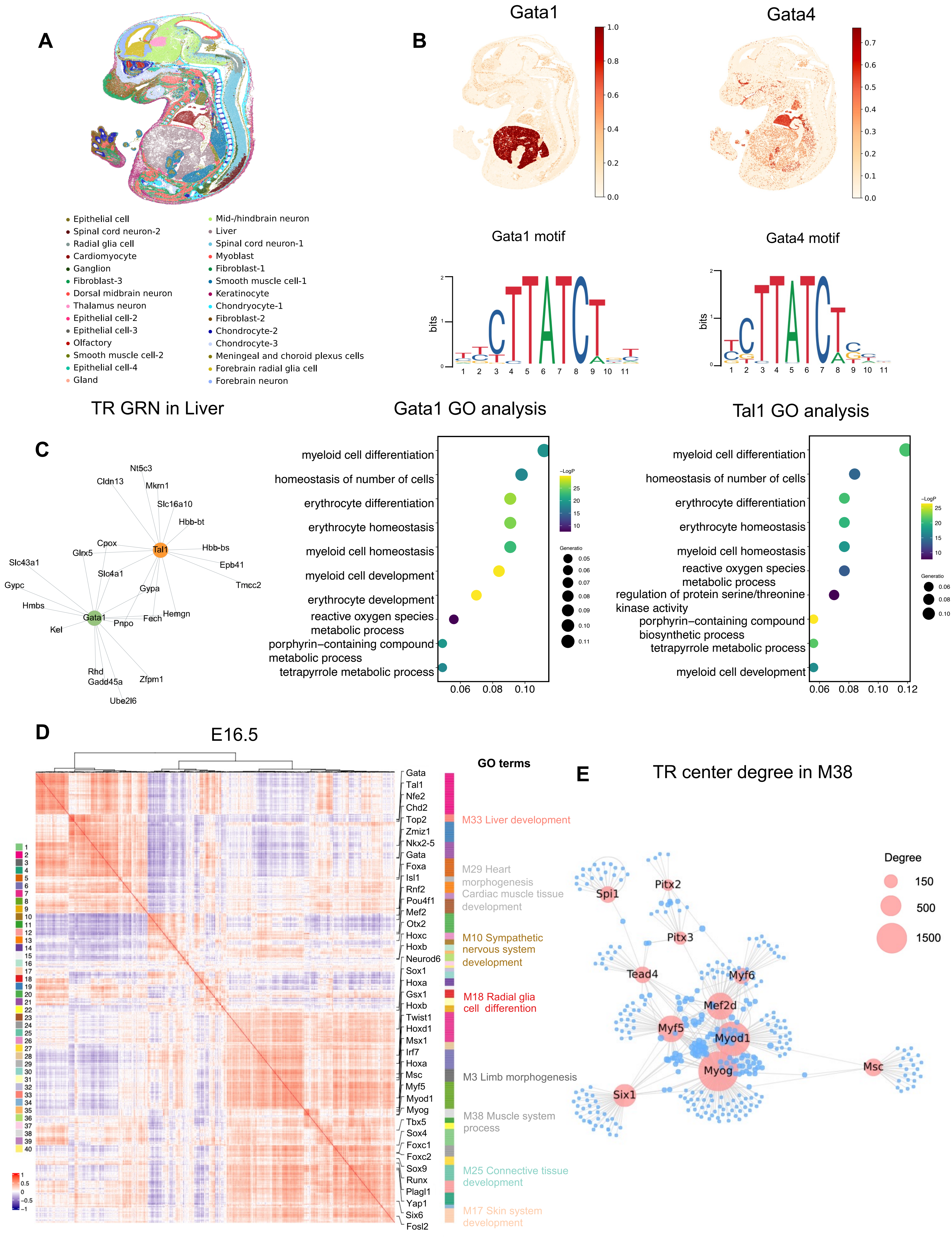

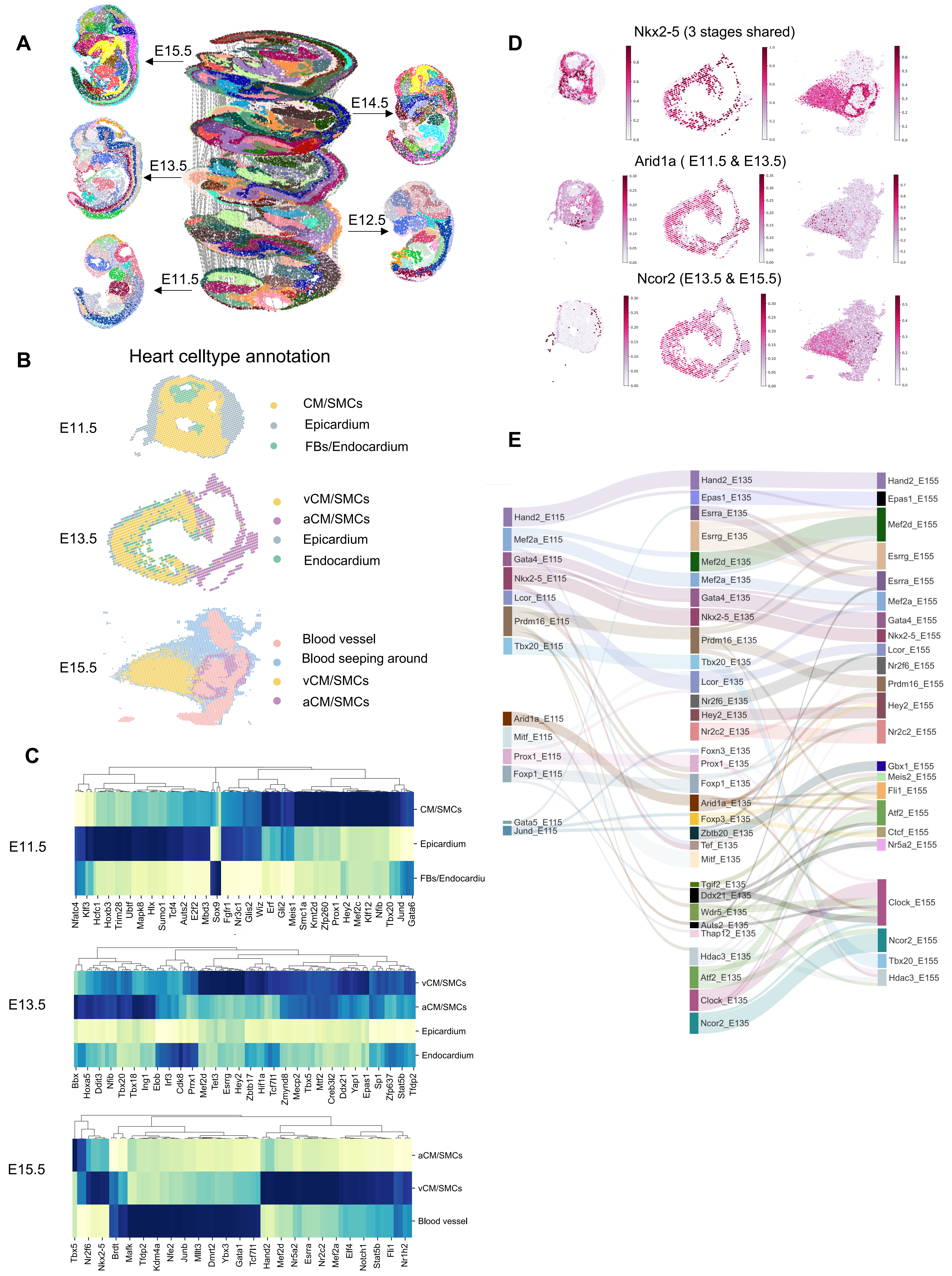

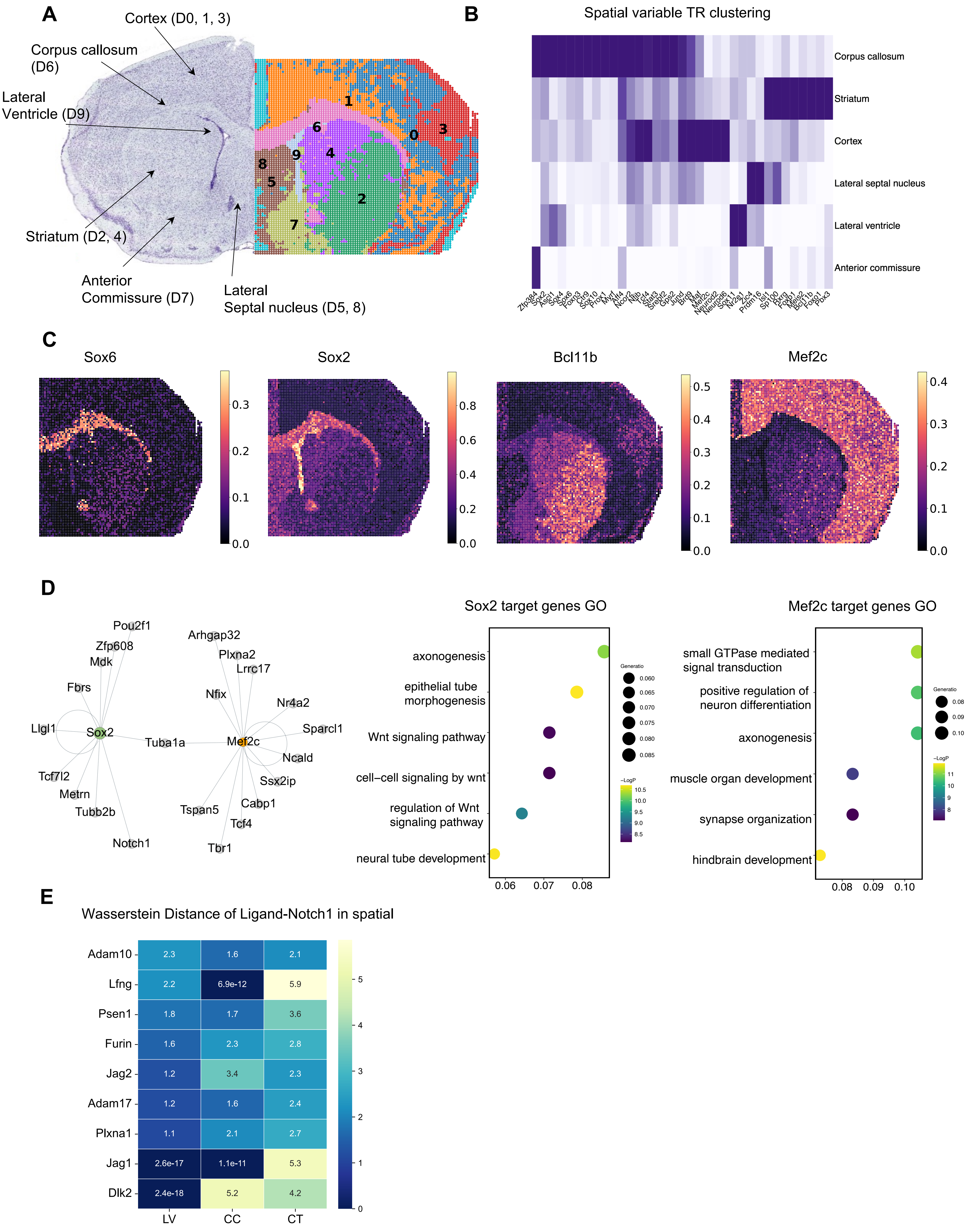
